# Sparse Modeling of Genomic Landscape Identifies Pathogenic Processes and Therapeutic Targets in Metastatic Breast Cancer

**DOI:** 10.1101/2023.08.29.554992

**Authors:** Mengchen Pu, Kai Tian, Weisheng Zheng, Xiaorong Li, Keyue Fan, Liang Zheng, Jielong Zhou, Yingsheng Zhang

**Affiliations:** StoneWise, AI, Ltd., Beijing, China; Minzu University of China, Beijing, China; Capital Normal University, Beijing, China

## Abstract

Breast cancer is a heterogeneous disease and ranks as one of the most lethal and frequently detected disease in the world. It poses significant challenges for precision therapy. To better decipher the patterns of heterogeneous nature in human genome and converge them into common functionals, mutational signatures are introduced to define the types of DNA damage, repair and replicative mechanisms that shape the genomic landscape of each cancer patient.

In this study, we developed a deep learning (DL) model, MetaWise 2.0, based on pruning technology that improved model generalization with deep sparsity. We applied it to patient samples from multiple sequencing studies, and identified statistically significant mutational signatures associated with metastatic progression using Shapley additive explanations (SHAP). We also employed gene cumulative contribution abundance analysis to link the mutational signatures with relevant genes, which could unearth the shared molecular mechanisms behind tumorigenesis and metastasis of each patient and lead to novel therapeutic target identification.

Our study illustrates that MetaWise 2.0 is an effective DL tool for discovering clinically meaningful mutational signatures in metastatic breast cancer (MBC) and relating them directly to relevant biological functions and gene targets. These findings could facilitate the development of novel therapeutic strategies and improve the clinical outcomes for individual patients.

## Introduction

Cancer genome are prone to numerous mutations and rearrangements that manifest genomic instability and heterogeneity. These variants modulate the expression and function of genes that regulate cell growth, differentiation, survival and migration^1,2^. The predominant cause of cancer-related morbidity and mortality is the metastatic spread, in which cancer cells disseminate from their primary site to other parts of the body through blood or lymphatic vessels. Metastatic processes typically involve cellular stressors and environmental shocks that elicit dramatic changes in the genome of cancer cells. These changes can bestow adaptive advantages to the cancer cells, such as enhanced invasiveness and therapeutic resistance. Therefore, elucidating the genomic characterization that underpins metastasis is crucial for devising effective strategies to prevent and treat cancers.

Breast cancer is the most common malignancy among women worldwide. For both primary and MBC, cumulative evidences point to the identification of the heterogeneous repertoire of disease-causing genes from various mutational processes^3,4^. However, due to the factors such as genomic background, lifestyle, tumor evolution and treatment pressure, the genomic alterations in metastatic breast cancer can differ drastically among patients. Thus, thorough genomic characterization of metastases for each patient will provide valuable insights, and is essential to understand the effects of systemic treatment on the tumor genome and improve the precision treatment of patients with metastatic breast cancers.

Genomic mutations can be characterized by mutational signatures defined as the proportion of mutations falling into mutational processes defined by their nucleotide context. Somatic mutations in whole genomes have empowered the detection of multiple mutational processes active in tumorigenesis, and such processes manifested in the tumor genome during tumor progression and treatment. Several studies have investigated the genomic landscape of metastatic tumors varieties^5-8^. Deep learning methods have been increasingly applied in the field of cancer research, particularly in studying the cancer metastasis progress. AI-based prediction models have been developed based on clinical data, such as medical images, gene expression profiles, *etc*^9-14^. At the same time, deep learning architectures are also being developed for the prediction of metastasis, including multilayer perceptron, convolutional neural networks, autoencoders, *etc*^15-18^. These models aim to solve a binary classification problem by classifying samples as either metastatic or non-metastatic. Several studies have shown the importance of mutational signatures in identifying cancer-associated genes and demonstrated some promising results on how these models guide precision medicine approaches^17,19^. However, major challenges remain to be addressed, including limited data accessibility, incorporation of multi-omics data, especially with extensive phenotypic annotations, limited generalizability across diverse patient cohorts, and the poor interpretability of DL models.

Over-parameterization, a common phenomenon in DL models, leads to high computational costs and impaired generalizability. Network pruning, the elimination of model components, has proven to be an effective technique that can optimize the efficiency of DL networks. This can help to reduce overfitting, improve model interpretability, and decrease computational requirements. In this study, we introduce an updated and extended version of MetaWise^17^. In comparison to our previous version, the new MetaWise 2.0 benefits from pruning technology, and offers model the generalization capability for data from different cohorts. Moreover, we integrated gene cumulative contribution abundance analysis with SHapley Additive exPlanations (SHAP) analysis to detect the significant correlations through association analysis of mutational signatures and key mutated functional genes and biological processes. These signatures were implicated in the growth of metastatic breast cancer cells, including those related to patient age, APOBEC enzymatic activity, DNA repair deficiency, *etc*. Several essential genes were identified to be the major biomarkers of MBC. Further enrichment analysis enabled the identification of various biological pathways involved in the development of MBC, such as the immune system processes, cell-cell communications, *etc*. The model also exhibited the capability to additionally stratify patients with MBC into more refined subgroups. This illustrates the potential utility of our updated approach, not only to discriminate the primary and metastatic cancers but also pinpoint the disease-associated molecular mechanisms for better therapeutic strategies of cancer treatment.

## Results

### Design of a sparse model for metastasis prediction

The diagram in Figure 1a illustrates the spread of MBC, originating from distinct primary sites and migrating toward several metastatic sites in our datasets. Apart from lymph nodes and breast as the locoregional metastatic sites, the most frequently observed distant metastatic organs in our dataset were liver, bone, skin and lung. As shown in Figure 1b, the distribution of molecular subtypes in our dataset differed between primary and metastatic breast cancer patients. In primary patients, ER+/HER- and triple-negative breast cancer (TNBC) were the two most common subtypes, accounting for 36.87% and 36.41% of cases respectively. In metastatic patients, ER+/HER2-remained the most common subtype but with more predominant compared to primary cases, accounting for 59.77% of metastatic cases. The remaining distribution of metastatic patients consisted of 12.63%, 9.59% and 3.98% TNBC, ER+/HER2+ and ER-/HER2+ respectively.

**Figure 1.**
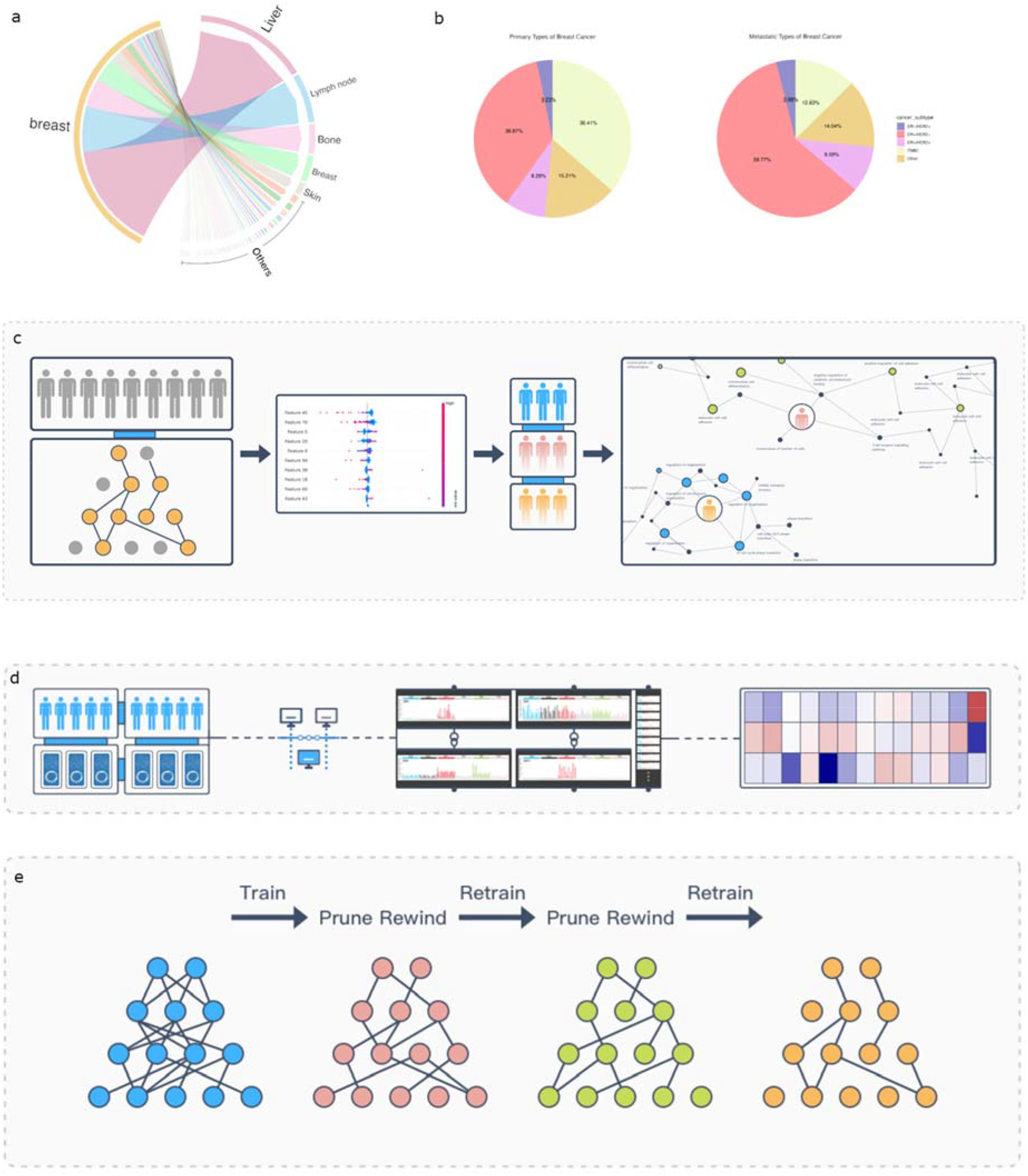
a. The Sankey diagram displays metastatic spreading directions from primary breast cancer toward several metastatic sites. Bandwidth is proportional to the number of metastatic tumor samples. Circle border thickness is proportional to the number of metastatic samples in that site. The color code representing the corresponding organ sites. b. The pie chart illustrates the subtypes distribution of primary and metastatic cancers within our dataset. The red, yellow, blue and grey color represents the ER+/HER2+, ER+/HER2-, ER-/HER2- and TNBC respectively. c. The workflow demonstrates our updated approach, MetaWise 2.0. d. The illustration of data pre-processing. e. The illustration of model pruning strategies.

As shown in Figure 1c, the workflow of our proposed model is illustrated, including a pruned deep learning predictor, an explanation module, and a sub-group gene enrichment analysis module. The predictor benefits from automated machine learning and model pruning techniques. To construct an optimal neural network architecture for predicting metastasis, we first utilized Bayesian optimization^20^, an efficient tool for hyperparameter tuning that employs a probabilistic model to guide the search through the configuration space towards the globally optimal design. After conducting an automated architecture search to identify the ideal configurations for architectural aspects such as the number of hidden layers and nodes per layer, as well as fine-tuning the hyperparameters, we applied pruning to the model to optimize it in two stages: input feature pruning and global weight pruning. Input feature pruning seeks to decrease the dimensionality of the input data by eliminating any irrelevant or redundant features. Global weight pruning takes a broader approach by removing weights from layers across the full model architecture. We used the magnitude-based criterion to rank the weights according to their absolute values, then removed a fixed percentage of weights with the smallest magnitudes, starting from 10% and increasing by 10% until 90%. We fine-tuned the sparse model for several epochs after each pruning step as shown in Figure 1e.

### Leveraging from Sparse model improves the performance of MetaWise-BC

In comparison to the model prior to the implementation of pruning techniques, it has been observed that the performance of the model post-pruning exhibits a marginal improvement with respect to the validation and internal test data. Furthermore, the model with pruned input features only or with pruned weights only showed slightly better performance in validation dataset than pruned input features and weights simultaneously, as shown in Table 1. This suggests that the pruning process may have facilitated the optimization of the model, enabling it to more effectively learn from feature correlation and even make slightly better predictions based on the training data with re-training after each pruning step.

**Table 1.** The average training and validation performance of the model across five folds of cross-validation.

| pruning | Val acc | Test acc | Test recall | Test precision | F1 score | Test AUC | Test AUPR |
| --- | --- | --- | --- | --- | --- | --- | --- |
| No | 91.5% | 86.9% | 90.2% | 94.3% | 92.2% | 78.4% | 96.5% |
| Weight pruning | 90.6% | 87.9% | 90.2% | 95.4% | 92.7% | 81.8% | 97% |
| Input pruning | 91.5% | 88.8% | 92.4% | 94.4% | 93.4% | 79.5% | 96.7% |
| Weight + input pruning | 91.5% | 86% | 89.1% | 94.3% | 91.6% | 78.0% | 96.4% |

Upon application to an external test data, it was observed that the three distinct pruning models exhibited substantial improvements in performance compared to their pre-pruning counterparts. Notably, there was a marked increase in recall, with an enhancement of 12%, while the F1 score demonstrated a 4.9% increase, and test accuracy and AUC both improved by 3.6%, as shown in Table 2. Additionally, other metrics also improved in varying degrees. These results convincingly demonstrate the power of pruning techniques to reduce model parameters, and help to mitigate overfitting. Also, the incorporation of pruning allows the model to better capture the key feature correlations, generalize to new data cohorts and accurately predict a patient’s potential risk of metastasis.

**Table 2.** The average performance on external data.

| pruning | Test acc | Test recall | Test precision | F1 score | Test AUC | Test AUPR |
| --- | --- | --- | --- | --- | --- | --- |
| No | 82.6 | 77.6 | 86.4 | 81.7 | 82.6 | 87.6 |
| Weight | 85.5 | 86.9 | 84.7 | 85.7 | 85.5 | 89.1 |
| Input | 83.8 | 82.2 | 84.9 | 83.5 | 83.8 | 88 |
| Weight + input | 85.5 | 86 | 85.2 | 85.6 | 85.5 | 89.1 |

### SHAP analysis interprets the model

To investigate the role of mutational signatures in breast cancer metastasis, we performed SHAP analysis on our model and identified certain signatures, such as SBS40, SBS1, SBS39, SBS8, SBS44, SBS2, SBS31 and three de novo signatures SBS_denovo_2, ID_denovo_3 and ID_denovo_4 which are involved in the development of metastatic breast cancers, as shown in Supplementary Figure 1. In the pruned and unpruned models, SHAP analysis revealed nearly the identical ten most impactful signatures. The key factors were consistent across both models, only with varied relative importance. The clock-like signature SBS1 is attributed to endogenous deamination of 5-methylcytosine to thymine^21^ which is related to age, as well as SBS40, which has also been shown to correlate with patient age in different types of human cancer^22^. The mutational signature SBS8 is common in most cancers, but its etiology is controversial. Recent evidence suggests that the SBS8 signature is due to DNA damage caused by late replication errors^23^, similar with defective DNA mismatch repair related signature SBS44. The uncharacterized signature SBS39 was significantly enriched in the basal subtype compared with three other breast cancer subtypes defined by PAM50 (Her2, Luminal A, Luminal B) ^24^. The SBS2 is associated with activity of the AID/APOBEC family of cytidine deaminases on the basis of similarities in the sequence context of cytosine mutations caused by APOBEC enzymes. The SBS31 is attributed to chemotherapy treatment with platinum drugs.

In order to examine the consistency between pruning strategies and SHAP analysis, we conducted a very radical pruning experiment by trimming 50% of the input features. The results showed that the eliminated features were consistently ranked lower in SHAP analysis with a Shapley value of 0.0 as shown in Supplementary Table 1. This suggests that the pruned input features had minimal impact on prediction outcomes, as determined by the interpretable analysis. Our findings indicate a high degree of consistency between the pruning strategy and SHAP analysis results, providing valuable insights into the effectiveness of these methods in improving model performance.

**Supplementary Table 1.** The Shapley value of removed features by pruning.

| Mutation signature | Shapley value |
| --- | --- |
| SBS6 | 0.0 |
| SBS7a | 0.0 |
| SBS7b | 0.0 |
| SBS12 | 0.0 |
| SBS16 | 0.0 |
| SBS17a | 0.0 |
| SBS19 | 0.0 |
| SBS21 | 0.0 |
| SBS22 | 0.0 |
| SBS25 | 0.0 |
| SBS26 | 0.0 |
| SBS28 | 0.0 |
| SBS30 | 0.0 |
| SBS33 | 0.0 |
| SBS35 | 0.0 |
| SBS36 | 0.0 |
| SBS38 | 0.0 |
| SBS41 | 0.0 |
| SBS88 | 0.0 |
| SBS92 | 0.0 |
| SBS93 | 0.0 |
| SBS_denovo_5 | 0.0 |
| SBS_denovo_6 | 0.0 |
| SBS_denovo_8 | 0.0 |
| SBS_denovo_9 | 0.0 |
| SBS_denovo_10 | 0.0 |
| SBS_denovo_11 | 0.0 |
| SBS_denovo_12 | 0.0 |
| SBS_denovo_13 | 0.0 |
| SBS_denovo_14 | 0.0 |
| SBS_denovo_15 | 0.0 |
| SBS_denovo_16 | 0.0 |
| SBS_denovo_20 | 0.0 |
| SBS_denovo_21 | 0.0 |
| SBS_denovo_22 | 0.0 |
| SBS_denovo_23 | 0.0 |
| SBS_denovo_24 | 0.0 |
| SBS_denovo_25 | 0.0 |
| SBS_denovo_26 | 0.0 |
| SBS_denovo_27 | 0.0 |
| DBS4 | 0.0 |
| DBS5 | 0.0 |
| DBS_denovo_5 | 0.0 |
| DBS_denovo_7 | 0.0 |
| ID3 | 0.0 |
| ID_denovo_2 | 0.0 |
| ID_denovo_5 | 0.0 |
| ID_denovo_6 | 0.0 |
| ID_denovo_7 | 0.0 |

**Supplementary Figure 1.**
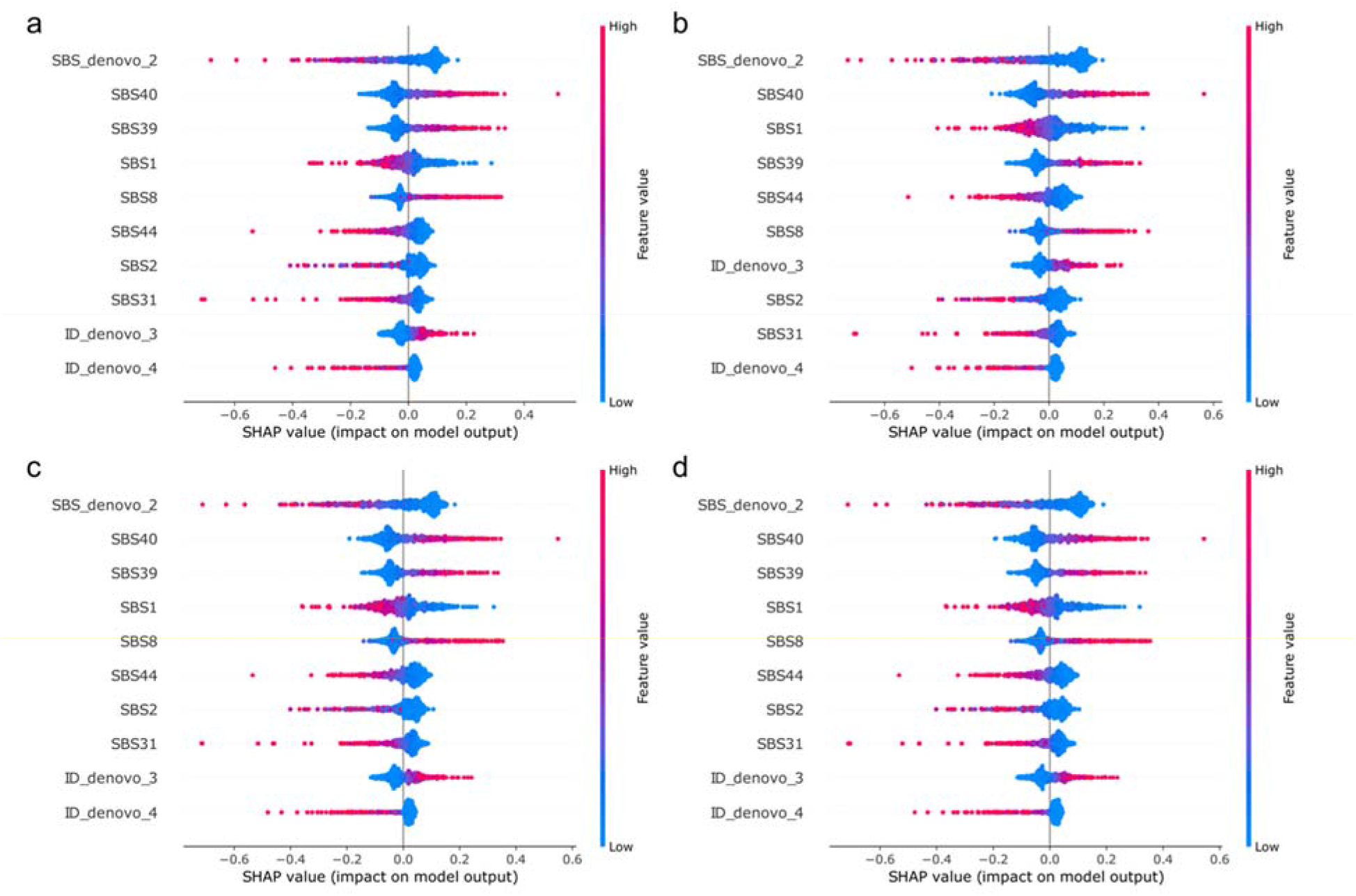
SHAP result of a. original model, b. Global weights pruned model, c. Input feature pruned model and, d. Global weights pruned and input pruned model.

To further validate the mutational signatures on model performance and results of the SHAP analysis, we conducted ablation studies. By systematically removing input mutation signatures with high Shapley value, we were able to assess the impact of these mutation signatures on model performance and understand the relationship between individual input mutation signatures and their contribution to prediction outcomes. The results of these ablation studies on top five features with large Shapley values are presented in Table 3. Our ablation studies support the effectiveness of SHAP analysis in identifying important mutational signatures and guiding the development of more accurate and interpretable models. After removing the top input features with the largest Shapley values, the models performance dropped significantly by approximately 5-10% on test accuracy. This suggests that the top features were critical to the model’s ability to predict. The sharp drop in performance after their removal highlights the importance of these features in the model and underscores the need for careful feature selection when developing predictive models.

**Table 3.** The model accuracy results from ablating the top features.

| Removed mutation signatures | Original | Weights | Input features | Weights and input features |
| --- | --- | --- | --- | --- |
| None | **0.831** | **0.841** | **0.827** | **0.850** |
| SBS_denovo_2 | 0.803 | 0.822 | 0.817 | 0.822 |
| SBS40 | 0.803 | 0.827 | 0.794 | 0.813 |
| SBS1 | 0.757 | 0.785 | 0.771 | 0.752 |
| SBS39 | 0.775 | 0.813 | 0.813 | 0.785 |
| SBS8 | 0.738 | 0.761 | 0.752 | 0.742 |

### Cumulative contribution abundance analysis and GO enrichment analysis reveals potentially relevant features

We conducted a comprehensive analysis to investigate the molecular mechanism behind mutation signatures by employing the RNMF method^25^ and gene enrichment analysis. A key component of this approach is the Cumulative Contribution Abundance (CCA) model, which effectively elucidates the associations between mutational signatures and genes. By calculating the cumulative contribution of each gene to each mutation signature, we were able to identify the genes that exerted the greatest influence on each mutation signature. This information was then applied to determine the genes that contributed most significantly to the mutation signature with the greatest impact for each sample. We subsequently grouped a subset of patients exhibiting similar characteristics of the most influential mutation signatures and obtained the most contributing genes corresponding to each patient within this subset. The resulting gene set was undertaken to gene enrichment analysis to further elucidate the underlying biological mechanisms.

Firstly, comparative analyses of the predicted genes enriched in primary and metastatic breast cancer samples were performed using KEGG, Reactome, and Gene Ontology databases (figure 2a). During the past decades, the molecular principles of metastasis remain an enigma even with the acceleration of multiomics research^26,27^. Genetic immune escape (GIE)^28^, microenvironment-derived epithelial to mesenchymal transition (EMT), cell motility^29,30^, breast cancer stem cells’ escape and sub localization are well characterized in the metastatic processes. In our study, the genes enriched in primary breast tumors are most correlated with cell growth and proliferation, cell homeostasis, and metabolism. Mutations of these genes can promote rapid cell proliferation, inhibit apoptosis, and ultimately lead to tumor formation. Nevertheless, the gene enriched in MBC are most correlated with immune system processes, cell communication, cell death *etc*. This implies these genes may participate in tumor metastasis and late-stage progression. The complement of these two sets showed significant differences in their corresponding biological functions. In the primary tumor enriched gene set, PI3K/AKT signaling, RAS-MAPK signaling, and ERBB signaling pathways were indicators of cell survival and proliferation; p53 signaling pathway, G1/S transition, and apoptotic response will affect cell death and cell cycle; carbon and lipids metabolism abnormality were associated with microenvironments, such as hypoxia and oxidative stress^31^. In comparison, four distinct pathways, including CCR3 pathway in inflammatory responses, PKC pathway, G protein signaling pathways and integration of energy metabolism, were specific to the metastasis enriched gene set. These pathways are associated with immune system processes, cell transformation and invasion, *etc*^32^.

**Figure 2.**
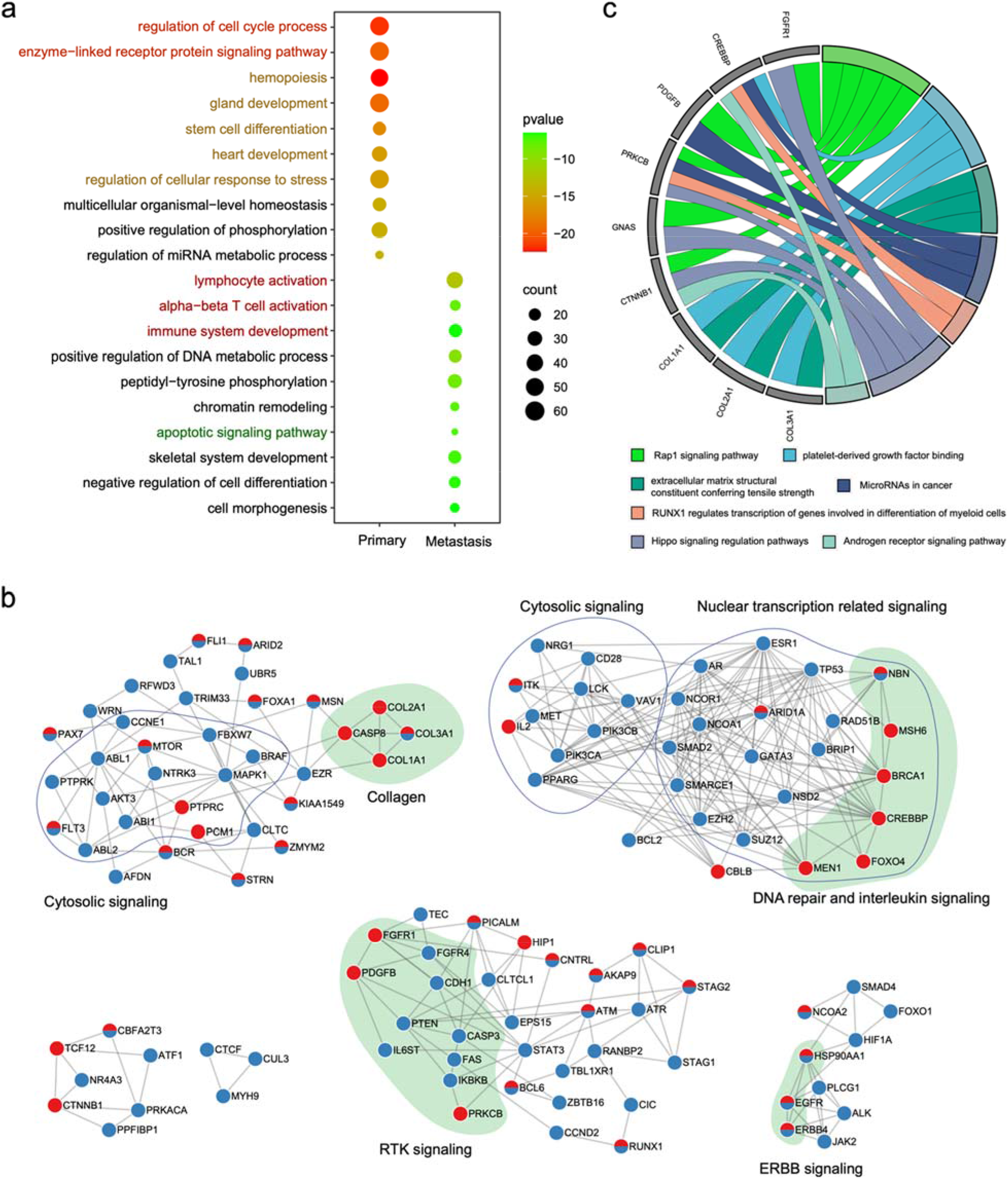
Visualizations of meta-analysis results of primary and metastasis gene sets. a. The gene ontology analysis of primary and metastasis enriched gene sets. b. The PPIN analysis of primary and metastasis enriched gene sets. c. The cellular processes analysis of nine metastasis specific genes.

To further decipher the functions of these two distinct gene sets, we integrated them into a protein-protein physical interaction network (PPIN) and filtered the interactions with a confidence score of less than 2.

The resulting genes were divided into six groups. As shown in Figure 2b, among these groups, five groups contained both primary and metastatic tumor genes, while the other group contained only primary tumor genes. The first group refers to cellular response to stimuli, especially immune response. The primary genes were mainly involved in RTK signaling and transcription regulation. In contrast, the metastasis genes were mainly involved in cellular surface communication and collagen recognition. The second group refers to transcription regulation. The primary tumor gene set was enriched in P53 signaling and hormone stimuli, while the metastasis genes were enriched in DNA repair and interleukin signaling. The other three groups of metastasis gene set were mainly enriched in Wnt signaling, RAP1 signaling, and calcium signaling pathway. The metastasis-specific genes were re-analyzed to characterize their functions in cellular processes. Nine genes, including growth factors, tyrosine kinases, GTPases, transcription factors and collagens, were involved in oncogenesis pathway (Figure 2c). For example, a PDGF gradient will drive cells to migrate towards the high concentration edge^33^; the extracellular matrix could be remodeled by different collagen types and concentrations to create a microenvironment supporting metastatic dissemination^34^.

To validate the potential of our model to effectively cluster patients, we analyzed the distinct groups within the metastatic patients. Eight clusters, including ID signature and SBS signature, were compared to elucidate their functional divergence. 3/56 genes were assigned to two clusters, and some genes in different clusters play the same function (Figure 3a). The top 20 GO terms showed significant differences between different clusters, e.g., SBS1 is mainly enriched in miRNA transcription, while SBS2 is involved in protein-containing complex localization. Genes enriched in different clusters function in different biological processes (Figure 3b).

**Figure 3.**
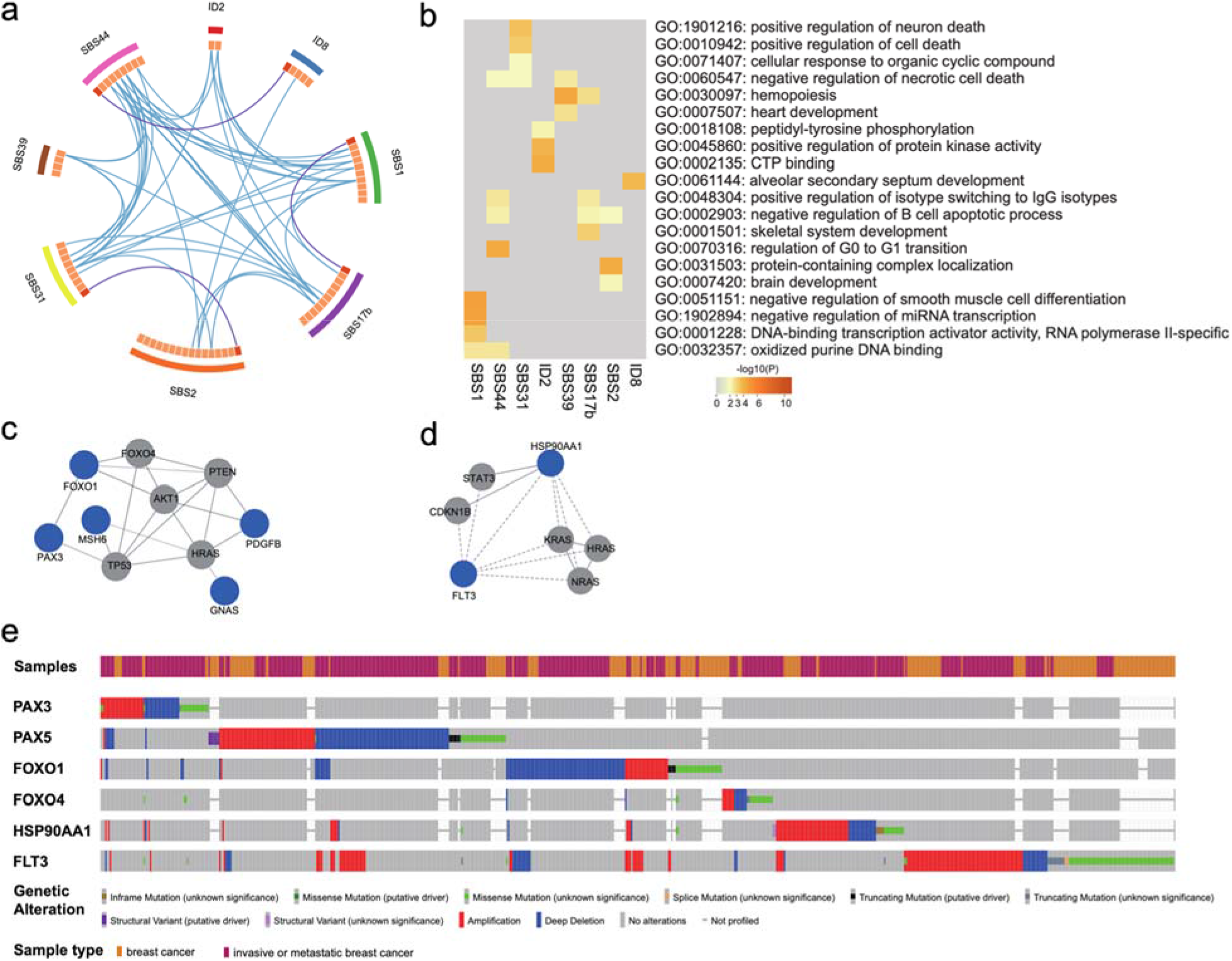
Visualizations of meta-analysis results based on multiple gene lists. A. predicted genes clustered well of sub-groups of metastatic patients. B. Heatmap showing the top enrichment clusters, one row per cluster, using a discrete color scale to represent statistical significance. Gray color indicates a lack of significance. C. The PPIN network of SBS1, the solid line represents the physical interactions which indicates that the proteins are part of a physical complex. The blue node represents the associated genes from mutational signatures. D. the PPIN network of ID1, the dot line represents the functional protein associations. E. Genetic alterations analysis of breast cancer with FOXO1, FOXO4, HSP90AA1 and FLT3.

To better understand the mechanisms that differentiate the subgroups, we selected two distinct groups with the different highest impact mutation signatures: ID2 and SBS1. Genetic alterations analysis and PPIN analysis were performed for both sub-groups, with the results presented in Figure 3. Our findings revealed that the gene sets of these two patient subgroups were enriched in entirely distinct molecular functions. The SBS1 signature indicates a C or T substitution and is believed to result from different forms of DNA damage^35^. Ten genes were enriched in our prediction category. FOXO4, PAX3, PLAG1 and NFIB are transcription factors; PDGFB is an RTK-related signaling protein; GNAS is a G-protein downstream of GPCR; VTI1A and SNX29 are related to protein sorting; and MSH6 is a DNA-binding protein which facilitates DNA mismatch repair. The functional process of FOXO4, PAX3, GNAS and PDGFB would be linked to the AKT pathway (Figure 3c). PAX5, a paralog of PAX3, has been validated to induce the gene expression of E-cadherin^36^ and MiR-215^37^ to inhibit the expression of FAK^38^, thereby suppressing breast cancer cell migration and invasion. A study of 263 breast cancer patients on cBioportal showed that 20 single missense mutations of PAX3 were related to metastatic breast cancer. Similarly, FOXO1 silencing in hepatocellular carcinoma causes ZEB2 expression and the EMT process^39^, but high expression of FOXO1 and FOXO3 upregulates matrix metalloproteinase (MMP) expression and enhances cancer cell metastasis^40,41^. Missense mutations of FOXO1 or FOXO4 were also correlated with breast cancer metastasis according to genetic alterations analysis based on cBioportal database (Figure 3e).

ID1 and ID2 were the result of replication slippage with the most happening of A or T indels at long poly(dA:dT) tracts^42-44^. The predicted ID2 cluster including FLT3 and HSP90AA1 have functions in protein kinase activity and CTP binding. HSP90 functioned as protein chaperone might have its dual character in breast cancer oncogenesis: decreased HSP90 has documented to proceed invasion and metastasis, whilst increased HSP90 enhances cell proliferation^45^. Intriguingly, HSP90AA1 was confirmed to be secreted extracellularly, and could activate EMT and migration^46,47^ (Figure 3d). These results indicate that our approach can successfully group patients in a way that reveals subsets enriched for distinct biological processes.

## Discussion

Over-parameterization is a common property in deep learning models, leading to increased computational costs and reduced generalization. As a remedy, network pruning has proven to be an effective technique to improve the efficiency of DL networks in situations where generalization is a concern and the computational cost is limited^48^. It is demonstrated in our work that pruning can be applied to the input layer of a neural network by removing input features with little to no impact on the output of the model. Pruning can simplify the model and improve its performance by reducing noise and focusing on the most relevant input features. Our results provide compelling evidence that our DL architecture is benefit from pruning technology. After pruning, the SHAP analysis and external test results indicate that pruned sparse model maintains the key input features and improves its generalization.

Our sparse model is capable of accurately identifying patients with different mutational profiles, which have significant implications for clinical management. Prior analysis methods can only associate mutational signatures with crude etiology. Via incorporating SHAP and gene accumulation analyses, our model not only can accurately predict metastasis risk in cancer patients, but for those who have metastasized, our approach can further stratify patients based on detailed molecular characterization with precision, portending favorable implications to inform personalized clinical management regimens. The capacity to segregate patients based on subtle genotypic distinctions enables tailored therapeutic interventions and prognostic predictions correlating with specific mutational profiles.

Further investigation will be justified to evaluate the clinical utility of this model, such as identifying actionable information from defined patient risk groups, identifying the significant relationship among mutations in coding and non-coding regions for metastasis. Due to the data confidentiality limitations, we did not conduct further research in this direction. It is important to investigate the interpretability of the features used by the model to predict prognosis for clinical guidance, which will be a topic of future work. In addition, having paired genomic data of the primary tumor and metastatic lesions from the same patients provides a powerful resource to study the genomic evolution of metastasis. Tracking the genomic changes from primary to metastatic sites in the same individual captures the trajectory of tumor evolution and progression in an authentic biological context, which will reveal core genomic features that distinguish metastatic clones from primary ones while eliminating the individual difference effects. Moreover, Martínez-Jiménez et. al. found that the metastatic tumors have a higher frequency of structural variants^8^. Incorporating additional structural mutation signature data and copy number variations (CNVs) has the potential to further improve AI algorithms from diverse multi-omics datasets. Leveraging diverse genomic data types, including structural variants, expression data across diverse populations worldwide will promote advances in AI for genomic medicines and enable more personalized therapies.

## Methods

### Genomic Data acquisition

The mutational signature contribution matrices pertaining to primary breast cancers within the Pan-Cancer Analysis of Whole Genomes (PCAWG) cohort, and metastatic breast cancers within the Hartwig Medical Foundation (HMF) cohort, were extracted from the supplementary tables of the reference^8^. The dataset encompasses two types of contribution matrices. The first one is denoted as “signature contributions”, encompassing the contributions for the mutational signatures detected in the two cohorts individually. The second type, termed “etiology contributions”, amalgamates the contributions of identical etiologies. For instance, contributions from mutational signatures SBS2 and SBS13 were conjoined to signify the collective contribution of the APOBEC etiology. In this study, we applied the “signature contributions” matrices for the training and validation of our models.

To constitute an external test dataset, we retrieved somatic mutation data from the BRCA-EU cohort, encompassing primary breast cancers, via the International Cancer Genome Consortium (ICGC) data portal (https://dcc.icgc.org/). Furthermore, we sourced somatic mutation data from the POG570 cohort^49^, containing metastatic breast cancers, from https://www.bcgsc.ca/downloads/POG570. In a bid to uphold data quality, samples characterized by low mutation burdens (mutation count < 50) were systematically excluded from the analysis. This curation process led to a refined dataset consist of 496 primary breast cancer samples and 127 metastatic breast cancer samples.

### Mutational signatures extraction

As shown in Figure 1d, to ascertain the mutational signature contributions from an external dataset, we implemented a comparable mutational signatures analysis pipeline, as delineated in reference^8^, albeit with certain modifications. In essence, we categorized somatic mutations within the external dataset into 96 SBS, 78 DBS, and 83 ID classes, as expounded upon in the earlier study^43^. The relative frequencies of each category within these channels were computed through utilization of the R package mutSigExtractor (https://github.com/UMCUGenetics/mutSigExtractor, v1.23). In order to ensure agreement with the mutational signatures employed in our model training and to mitigate potential bleeding effects stemming from the mutational signature extraction process, we refrained from conducting de novo mutational signatures extraction on the external dataset. Instead, we opted to leverage the 14 SBS, 5 DBS, and 9 ID mutational signatures previously identified within breast cancer instances within the PCAWG and HMF cohorts, employing these as reference signatures. Then, the fitToSignatures() function of mutSigExtractor, which employing a least square fitting algorithm, was applied to ascertain the individualized contributions of the aforementioned reference signatures within each sample from the external dataset. The matrices denoting these contribution values were subsequently employed as input features for evaluating the performance of our model.

### Implementation of MetaWise 2.0

The updated MetaWise framework consists of two modules: the classification module and the pathogenic process identification module. The workflow is shown is Figure 1c. The classification module aims to predict the metastasis possibility of a patient based on their genomic data. The pathogenic process identification module aims to identify the key biological processes that are associated with metastasis process from the genomic profile of patients.

#### Model design and evaluation

We implemented our model using the Keras framework with Tensorflow as the backend. Our model consists of fully-connected layers, each followed by a batch normalization layer and a ReLU activation function, with a softmax output layer. To optimize the performance of our model, we fine-tuned various hyperparameters, such as the learning rate, the weight decay, the dropout rate, and the activation function. The pruning process was performed by Keras prune_low_magnitude function.

We evaluated the performance of the updated approach and compared with our previous model, MetaWise^17^ by five-fold cross validation. We measured the accuracy, recall, precision, specificity, F1-score, the matthews correlation coefficient (MCC) and the area under the receiver operating characteristic curve (AUROC) and the area under the precision recall curve (AUPR) on both internal and external test sets. We compared the results of weight pruning, input feature pruning and global pruning to identify the best pruning strategy.

#### Model interpretability analysis

We employed SHAP (SHapley Additive exPlanations) analysis to gain insights into the explanation of our model and to reveal the significance and impact of various mutation signatures on the prediction. SHAP analysis can offer both global and local explanations of the model, as well as feature interactions and dependencies. We utilized the Python library, shap^50^, to conduct SHAP analysis for our model. After we identified the important mutation signatures for each sub-set data, we associated genes with mutational signatures using gene cumulative contribution abundance analysis^25^ to understand the pathogenesis of patients.

#### Cumulative contribution abundance analysis

In order to in-depth mine the relationship between genes and mutational signatures, the mutational signature analysis was performed to associate mutational signatures with genes. A simple and practical R package, RNMF^25^, was applied by analyzing cumulative contribution abundance of genes.

#### GO enrichment analysis

Gene Ontology (GO) term enrichment analysis was performed to identify over-represented GO terms in our gene set of interest. The GO system of classification assigns genes to a set of predefined bins based on their functional characteristics. Enrichment analysis identifies which GO terms are over-represented (or under-represented) in the gene set using annotations for that gene set. Enrichment analysis was performed using the clusterProfiler^51^, which is an R package that provides functions for statistical analysis and visualization of functional profiles for genes and gene clusters. It can be used to identify enriched gene sets in a cluster of genes, or to compare the functional profiles of two or more clusters. The GO aspect (molecular function, biological process, cellular component) for the analysis was selected, as well as the species from which the genes come. The results page displays a table that lists significant shared GO terms used to describe the set of genes entered.

### Statistical analysis and results visualization

#### Ablation experiment

To perform the ablation experiments, we first ranked the input features according to their Shapley values. Then, we removed the top-ranked input features one at a time, retrained the model on the remaining features, and evaluated its performance using a variety of metrics. The results of these ablation experiments were analyzed to determine the impact of each removed input feature on model performance.

#### meta-analysis

Additional pathway enrichment analyses, gene list annotations and protein-protein interaction network analysis were performed using the free online meta-analysis tool Metascape^52^ (https://metascape.org/).

## Code availability

Code will be uploaded to the github repository (https://github.com/promethiume/MetaWise) once the paper has been conditionally accepted, and are available from the corresponding author on reasonable request during the manuscript review process.

## Acknowledgements

We thank G. Peng, Y. Xin and L. Wei from the Innovation Center of StoneWise, AI. Ltd. for their helpful discussions and support; X.Xiang, S.Guo, Y.Wang and our colleagues at StoneWise, AI. Ltd. for their support and encouragement.

## Author contributions statement

Y.Z. and M.P. initiated the project. M.P., X.L. and K.F. conducted the research. W.Z. and K.F. curated and pre-processed the data. M.P. designed the method. M.P., X.L, K.F. and K.T. performed the evaluation and analyzed the results. L.Z. contributed on data visualization. M.P., and K.T. wrote the manuscript. All authors reviewed manuscript.

## Additional information

The authors declare no competing interests.

